# Brain dysfunction in chronic pain patients assessed by resting-state electroencephalography

**DOI:** 10.1101/595470

**Authors:** Son Ta Dinh, Moritz M. Nickel, Laura Tiemann, Elisabeth S. May, Henrik Heitmann, Vanessa D. Hohn, Günther Edenharter, Daniel Utpadel-Fischler, Thomas R. Tölle, Paul Sauseng, Joachim Gross, Markus Ploner

**Author notes:** **Corresponding author:** Markus Ploner, Department of Neurology, Technische Universität München, Ismaninger Str. 22, 81675 Munich, Germany, Fon.

## Abstract

Chronic pain is a common and severely disabling disease whose treatment is often unsatisfactory. Insights into the brain mechanisms of chronic pain promise to advance the understanding of the underlying pathophysiology and might help to develop disease markers and novel treatments. Here, we systematically and comprehensively exploited the potential of electroencephalography (EEG) to determine abnormalities of brain function during the resting state in chronic pain. To this end, we performed state-of-the-art analyses of oscillatory brain activity, brain connectivity and brain networks in 101 patients of either sex suffering from chronic pain. The results show that global and local measures of brain activity did not differ between chronic pain patients and a healthy control group. However, we observed significantly increased connectivity at theta (4 – 8 Hz) and gamma (> 60 Hz) frequencies in frontal brain areas as well as global network reorganization at gamma frequencies in chronic pain patients. Furthermore, a machine learning algorithm could differentiate between patients and healthy controls with an above-chance accuracy of 57%, mostly based on frontal connectivity. These results implicate increased theta and gamma synchrony in frontal brain areas in the pathophysiology of chronic pain. While substantial challenges concerning accuracy, specificity and validity of potential EEG-based disease markers remain to be overcome, our study identifies abnormal frontal synchrony at theta and gamma frequencies as promising targets for non-invasive brain stimulation and/or neurofeedback approaches.

## Introduction

Chronic pain is a disease characterized by ongoing pain and significant sensory, cognitive and affective abnormalities [55; 85] which have detrimental effects on quality of life. Affecting between 20 and 30 percent of the adult population [11; 40], chronic pain is a leading cause of disability worldwide [32]. Thus, advances in the understanding and treatment of chronic pain are urgently needed.

Studies in animals and humans have revealed that chronic pain is associated with significant structural and functional changes of the brain [4; 43; 62]. In particular, the prefrontal cortex and subcortical brain areas including amygdala, hippocampus and striatal areas have been implicated in chronic pain [4; 5; 35; 53; 62; 69; 80]. Further insights into the brain mechanisms of chronic pain not only promise to advance the understanding of the underlying pathophysiology but could also be clinically highly useful. In particular, a brain-based marker of chronic pain could be immensely helpful for the diagnosis, classification and treatment of pain [18; 79]. Accordingly, the feasibility, limitations and perspectives of brain-based markers of pain are currently intensively discussed in the pain research community [18; 56; 73] and beyond [49; 63; 94]. Recent functional magnetic resonance imaging studies have made important first steps towards such a brain-based marker of experimental [89] and chronic pain [50; 51].

Using electroencephalography (EEG) to assess abnormalities of brain function and to establish a brain-based marker of chronic pain is particularly appealing as it is safe, cost-effective, broadly available and potentially mobile. Moreover, an EEG-based marker of chronic pain might not only be helpful for the diagnosis and classification of chronic pain but might itself represent a target for novel therapeutic strategies such as neurofeedback [72] or non-invasive brain stimulation techniques [60]. As a potential first step in that direction, some EEG studies have shown a slowing of neural oscillations together with an increase of oscillatory brain activity at theta frequencies (4 – 8 Hz) in chronic pain patients [65; 84]. These observations have been embedded in the thalamocortical dysrhythmia model of neuropsychiatric disorders [47]. In this model, abnormal thalamocortical theta activity yields abnormal oscillations at gamma (> 30 Hz) frequencies which eventually result in different neuropsychiatric symptoms including ongoing pain. This model is highly appealing, but evidence is sparse, contradictory and mostly confined to small groups of patients suffering from neuropathic subtypes of chronic pain [58; 59]. Thus, a general EEG-based marker of chronic pain remains to be demonstrated.

In the present study, we aimed to systematically and comprehensively exploit the potential of EEG to determine abnormalities of resting-state brain activity as a potential brain-based marker of chronic pain. In a large cohort of chronic pain patients and age- and gender-matched healthy controls, we analyzed global and local measures of oscillatory brain activity. Moreover, we performed connectivity analyses using phase-based and amplitude-based measures in source space as well as graph theory-based network analyses. In a univariate approach, we statistically compared these measures between patients and healthy controls. Moreover, in a multivariate machine learning approach, we tested whether patterns of these measures allow to distinguish between chronic pain patients and healthy controls.

## Materials and Methods

### Participants

101 patients (age 58.2 ± 13.5 years (mean ± standard deviation), 69 female) suffering from chronic pain and 84 age- and gender-matched healthy control participants (age 57.8 ± 14.6 years, 55 female) participated in the study. Inclusion criteria for patients were a clinical diagnosis of chronic pain, with pain lasting at least 6 months and a minimum reported average pain intensity ≥ 4/10 during the last 4 weeks (0 = no pain, 10 = worst imaginable pain). Exclusion criteria were acute changes of the pain condition during the last 3 months, for example due to recent injuries or surgeries. Further exclusion criteria were major neurological diseases such as stroke, epilepsy or dementia, major psychiatric diseases aside from depression and severe general diseases. Finally, patients on medication with benzodiazepines were excluded, other medication was not restricted. Demographic and clinical details of participants are shown in Tables 1 and 2, respectively. All participants gave written informed consent. The study was approved by the ethics committee of the Medical Faculty of the Technische Universität München and carried out in accordance with the relevant guidelines and regulations. A power analysis for an independent two-sample t-tests using G*Power [25] showed that the number of participants allowed for detecting differences between groups of at least medium effect size (Cohen’s d = 0.5) with a statistical power of 0.9.

**Table 1.** Demographics and questionnaire results. CP, chronic pain patients; HC, healthy controls; VR-12, Veteran’s RAND 12; PCS, physical component score; MCS, mental component score; VAS, visual analog scale; Avg. pain intensity, average pain intensity in the last 4 weeks; PDI, pain disability index; PDQ, painDETECT.

| | CP (N = 101, mean ± sd) | HC (N = 84, mean ± sd) |
| --- | --- | --- |
| Gender (M/F) | 32/69 | 29/55 |
| Age | 58.2 ± 13.5 | 57.8 ± 14.6 |
| VR-12 PCS score | 31.7 ± 7.8 | 52.74 ± 5.7 |
| VR-12 MCS score | 46.4 ± 12.0 | 54.1 ± 9.0 |
| Current pain intensity (VAS) | 5.2 ± 1.9 | - |
| Avg. pain intensity (VAS) | 5.7 ± 1.6 | - |
| Pain duration (months) | 121.9 ± 114.3 | - |
| PDI score | 28.3 ± 14.7 | - |
| PDQ score | 17.4 ± 6.5 | - |

**Table 2.** Patient characteristics. ID, patient identification number; ys, years; Curr. pain, currently experienced pain; Avg. pain, average pain in the last 4 weeks; VAS, visual analog scale: 0 = no pain, 10 = worst imaginable pain; MQS, medication quantification scale; BDI, Beck Depression Inventory II, score ≥ 18 = clinically relevant depression; PDQ, painDETECT, score ≥ 19 = neuropathic pain component probable; m, male; f, female; CBP, chronic back pain; PNP, poly-neuropathic pain; NP, neuropathic pain, PHN, post-herpetic neuralgia; CWP, chronic widespread pain; JP, joint pain; n.a., not available because the respective questionnaire was not completed.

| ID | Age (ys) | Sex | Pain duration (months) | Curr. pain (VAS) | Avg. Pain (VAS) | Pain diagnosis | MQS | BDI | PDQ |
| --- | --- | --- | --- | --- | --- | --- | --- | --- | --- |
| 1 | 67 | m | 360 | 5 | 5 | CBP | 4 | 3 | 30 |
| 2 | 54 | f | 120 | 4 | 5 | CBP | 32 | 35 | 14 |
| 3 | 64 | f | 96 | 5 | 7 | CBP | 11 | 7 | 16 |
| 4 | 41 | m | 24 | 7 | 6 | CBP | 6 | 15 | 22 |
| 5 | 74 | m | 180 | 3 | 5 | CBP | 7 | 18 | 4 |
| 6 | 58 | f | 168 | 7 | 6 | CBP | 19 | 22 | 21 |
| 7 | 65 | m | 48 | 6 | 5 | CBP | 4 | 11 | 12 |
| 8 | 65 | f | 132 | 3 | 3 | CBP | 6 | 5 | 24 |
| 9 | 76 | f | 24 | 5 | 5 | CBP | 4 | 9 | 20 |
| 10 | 33 | f | 36 | 6 | 7 | CBP | 9 | 19 | 16 |
| 11 | 45 | f | 12 | 8 | 7 | CBP | 5 | 20 | 8 |
| 12 | 51 | f | 24 | 5 | 6 | CBP | 11 | 22 | 19 |
| 13 | 73 | f | 108 | 6 | 8 | CBP | 11 | 21 | 26 |
| 4 | 41 | f | 120 | 5 | 6 | CBP | 13 | 14 | 13 |
| 15 | 55 | f | 252 | 9 | 9 | CBP | 24 | 24 | 20 |
| 16 | 73 | m | 300 | 5 | 4 | CBP | 4 | 10 | 5 |
| 17 | 46 | m | 360 | 6 | 7 | CBP | 16 | 10 | 5 |
| 18 | 50 | m | 60 | 7 | 7 | CBP | 0 | 5 | 17 |
| 19 | 59 | f | 24 | 4 | 6 | CBP | 12 | 31 | 23 |
| 20 | 62 | m | 12 | 5.5 | 6 | CBP | 10 | 26 | 22 |
| 21 | 54 | m | 84 | 5 | 7 | CBP | 16 | 18 | 17 |
| 22 | 39 | f | 144 | 4 | 2 | CBP | 0 | 12 | 22 |
| 23 | 66 | m | 48 | 4 | 4 | CBP | 8 | 19 | 10 |
| 24 | 57 | m | 300 | 5 | 6 | CBP | 22 | 12 | 17 |
| 25 | 52 | m | 300 | 4 | 3 | CBP | 11 | 22 | 13 |
| 26 | 47 | f | 180 | 4 | 3.5 | CBP | 26 | 19 | 15 |
| 27 | 24 | f | 96 | 7.5 | 7.5 | CBP | 5 | 13 | 20 |
| 28 | 59 | f | 180 | 3 | 5 | CBP | 6 | 4 | 10 |
| 29 | 82 | m | 60 | 5 | 5 | CBP | 14 | 21 | 8 |
| 30 | 54 | m | 120 | 6.5 | 6 | CBP | 12 | 17 | 22 |
| 31 | 62 | f | 36 | 3 | 4.5 | CBP | 3 | 22 | 11 |
| 32 | 73 | m | 480 | 8 | 8 | CBP | 4 | 11 | 10 |
| 33 | 48 | m | 64 | 0.5 | 4 | CBP | 24.45 | 24 | 22 |
| 34 | 55 | f | 240 | 5 | 6 | CBP | 8.55 | 31 | 11 |
| 35 | 59 | f | 60 | 3 | 4 | PNP | 13.5 | 11 | 15 |
| 36 | 48 | f | 108 | 3 | 4 | PHN | 15.4 | 13 | 28 |
| 37 | 73 | f | 24 | 5 | 5 | CBP | 13.5 | 5 | 11 |
| 38 | 57 | m | 84 | 8 | 7.5 | PNP | 22.35 | 15 | 33 |
| 39 | 57 | f | 96 | 6 | 7 | CBP | 3.4 | 20 | 15 |
| 40 | 77 | f | 36 | 5 | 7.8 | PNP | 4 | 10 | 11 |
| 41 | 77 | f | 480 | 3.5 | 3.5 | CBP | 8.6 | 9 | 19 |
| 42 | 53 | f | 24 | 4 | 5 | CBP | 16.3 | 12 | 24 |
| 43 | 80 | m | 48 | 4 | 4 | PHN | 11.4 | 4 | 18 |
| 44 | 77 | f | 192 | 4 | 6 | CWP | 12.8 | 0 | 23 |
| 45 | 57 | f | 72 | 5 | 6 | CWP | 5.7 | 42 | 17 |
| 46 | 67 | f | 21 | 6 | 8 | NP | 3.4 | 8 | 17 |
| 47 | 77 | f | 324 | 6 | 6 | CWP | 3.4 | 16 | 24 |
| 48 | 61 | f | 36 | 4.5 | 6 | NP | 3.4 | 7 | 21.5 |
| 49 | 65 | f | 23 | 8 | 8 | CWP | 4 | 7.5 | 25 |
| 50 | 65 | f | 180 | 5 | 4 | CWP | 0 | 12 | 22 |
| 51 | 56 | f | 213 | 4 | 7 | CWP | 8 | 13 | 23 |
| 52 | 57 | m | 264 | 3 | 4 | JP | 13.25 | 10 | 11 |
| 53 | 41 | f | 24 | 4 | 5 | NP | 25.5 | 12 | 24 |
| 54 | 69 | f | 72 | 8 | 5 | CBP | 0 | 19 | 13 |
| 55 | 56 | f | 108 | 8 | 8 | PNP | 17.2 | 10 | 25 |
| 56 | 72 | f | 120 | 7 | 8 | CWP | 15.4 | 16 | 20 |
| 57 | 57 | m | 96 | 5 | 7 | PNP | 10.9 | 11 | 28 |
| 58 | 82 | f | 36 | 2 | 5 | CBP | 4.6 | n.a. | 10 |
| 59 | 70 | f | 420 | 4 | 6 | CWP | 14.8 | 28 | 22 |
| 60 | 70 | f | 72 | 4 | 6 | PHN | 10.3 | 11 | 22 |
| 61 | 54 | f | 24 | 7 | 5 | PHN | 5.8 | 10 | 10 |
| 62 | 69 | f | 48 | 6 | 5 | PHN | 8.4 | 6 | 16 |
| 63 | 66 | f | 84 | 2 | 3 | JP | 6 | 4 | 13 |
| 64 | 52 | f | 204 | 6 | 6 | CWP | 10.2 | 16 | 21 |
| 65 | 77 | f | 72 | 8 | 8 | JP | 4.6 | n.a. | 19 |
| 66 | 42 | m | 252 | 7 | 9 | CBP | 17.35 | 33 | 22 |
| 67 | 51 | f | 31 | 6 | 5 | JP | 9.1 | 15 | 7 |
| 68 | 55 | m | 7 | 7 | 7 | JP | 6.9 | 1 | 13 |
| 69 | 68 | f | 24 | 1 | 4 | NP | 3.8 | 0 | 23 |
| 70 | 71 | m | 324 | 4 | 5 | CBP | 10.8 | 5 | 6 |
| 71 | 24 | f | 108 | 6 | 6 | CWP | 8 | 31 | 21 |
| 72 | 71 | m | 36 | 6 | 6 | CBP | 5.8 | 9 | 14 |
| 73 | 68 | f | 204 | 3 | 4 | JP | 8.8 | 7 | 15 |
| 74 | 86 | f | 120 | 3 | 3 | CBP | 2 | 5 | 6 |
| 75 | 68 | f | 120 | 4 | 5 | NP | 7.8 | 5 | 28 |
| 76 | 45 | m | 48 | 1 | 2 | CBP | 3.8 | 10 | 9 |
| 77 | 18 | f | 16 | 4 | 6 | PHN | 0 | 14 | 20 |
| 78 | 80 | f | 25 | 8 | 7 | PHN | 23.1 | 15 | 20 |
| 79 | 60 | m | 60 | n.a. | n.a. | PNP | 0 | n.a. | n.a. |
| 80 | 60 | f | 54 | 3.5 | 5.5 | CBP | 13.5 | 22 | 18 |
| 81 | 57 | m | 17 | 6.5 | 6.5 | CBP | 24.3 | 25 | 24 |
| 82 | 45 | m | n.a. | 7.3 | n.a. | CWP | 0 | 17 | n.a. |
| 83 | 24 | f | n.a. | 3.2 | n.a. | CWP | 0 | 17 | n.a. |
| 84 | 49 | m | n.a. | 7.3 | n.a. | CWP | 0 | 39 | n.a. |
| 85 | 47 | f | n.a. | 2.4 | n.a. | CWP | 0 | 21 | n.a. |
| 86 | 53 | f | n.a. | 7.5 | n.a. | CWP | 0 | 22 | n.a. |
| 87 | 41 | f | n.a. | 6.2 | n.a. | CWP | 0 | 29 | n.a. |
| 88 | 46 | m | n.a. | 7.4 | n.a. | CWP | 0 | 20 | n.a. |
| 89 | 56 | f | n.a. | 1.8 | n.a. | CWP | 0 | 22 | n.a. |
| 90 | 55 | f | n.a. | 7.8 | n.a. | CWP | 0 | 30 | n.a. |
| 91 | 60 | f | n.a. | 6.2 | n.a. | CWP | 0 | 14 | n.a. |
| 92 | 55 | f | n.a. | 8.6 | n.a. | CWP | 0 | 25 | n.a. |
| 93 | 48 | f | n.a. | 2.9 | n.a. | CWP | 0 | 27 | n.a. |
| 94 | 59 | m | n.a. | 6.8 | n.a. | CWP | 0 | 13 | n.a. |
| 95 | 60 | f | n.a. | 5.0 | n.a. | CWP | 0 | 17 | n.a. |
| 96 | 58 | f | n.a. | 4.5 | n.a. | CWP | 0 | 12 | n.a. |
| 97 | 71 | f | n.a. | 8.4 | n.a. | CWP | 0 | 36 | n.a. |
| 98 | 42 | m | n.a. | 3.7 | n.a. | CWP | 0 | 12 | n.a. |
| 99 | 38 | f | n.a. | 6.5 | n.a. | CWP | 0 | 10 | n.a. |
| 100 | 65 | f | n.a. | 8.8 | n.a. | CWP | 0 | 18 | n.a. |
| 101 | 60 | f | n.a. | 6.0 | n.a. | CWP | 0 | 21 | n.a. |

### Recordings

Brain activity was recorded using EEG. Recordings were performed during the resting-state, i.e. participants were asked to stay in a relaxed and wakeful state, without any particular task. EEG data were recorded with eyes closed and eyes open for 5 minutes each. As the eyes closed condition showed better data quality and less muscle artifacts, analyses were focused on this condition.

EEG data were recorded using 64 electrodes consisting of all 10-20 system electrodes and the additional electrodes Fpz, CPz, POz, Oz, Iz, AF3/4, F5/6, FC1/2/3/4/5/6, FT7/8/9/10, C1/2/5/6, CP1/2/3/4/5/6, TP7/8/9/10, P5/6 and PO1/2/9/10 plus two electrodes below the outer canthus of each eye (Easycap, Herrsching, Germany) and BrainAmp MR plus amplifiers (Brain Products, Munich, Germany). All EEG electrodes were referenced to FCz and grounded at AFz. Simultaneously, muscle activity was recorded with two bipolar surface electromyography (EMG) electrode montages placed on the right masseter and neck (semispinalis capitis and splenius capitis) muscles [17] and a BrainAmp ExG MR amplifier (Brain Products, Munich, Germany). The EMG ground electrode was placed at vertebra C2. All data were sampled at 1000 Hz (0.1 μV resolution) and band-pass filtered between 0.016 Hz and 250 Hz. Impedances were kept below 20 kΩ.

Prior to the EEG recordings, patients completed the following questionnaires to assess pain characteristics and comorbidities: short-form McGill pain Questionnaire (SF-MPQ) [54], Pain Disability Index (PDI) [21], painDETECT (PDQ) [29], Beck Depression Inventory II (BDI-II) [8], State-Trait-Anxiety Inventory (STAI) [74] and the Veteran’s RAND 12-Item Health Survey (VR-12) [68].

### Preprocessing

Preprocessing was performed using the BrainVision Analyzer software (Brain Products, Munich, Germany). Data were downsampled to 250 Hz. For artifact identification, a high-pass filter of 1 Hz and a 50 Hz notch filter for line noise removal were applied to the EEG data. Independent component analysis was performed [39] and components representing eye movements and muscle artifacts were identified based on time courses and topographies. Furthermore, time intervals of 400 ms around data points with signal jumps exceeding ± 100 µV were marked for rejection. Lastly, all data were visually inspected and remaining bad segments marked. Subsequently, independent components representing artifacts were subtracted from the raw, unfiltered EEG data [93] and EEG data were re-referenced to the average reference. The reference electrode FCz was added to the signal array.

Next, data were exported from the BrainVision Analyzer and further analyzed in Matlab (Mathworks, Natick, MA, USA) with the FieldTrip [57] and Brain Connectivity Toolbox [64], along with custom-written code. Data were segmented into 2 s epochs with 1 s overlap. A 2 s epoch length was chosen to balance the stationarity of the signals and the number of samples for lower frequencies (down to 4 Hz) [14; 81]. All analyses are based on these epochs and the following four frequency bands: theta (4 – 8 Hz), alpha (8 – 13 Hz), beta (14 – 30 Hz) and gamma (60 – 100 Hz). We observed strong non-stationary line noise around 45 – 55 Hz and therefore excluded the “low gamma” band from frequency band specific analyses.

### EEG analysis – overview

Figure 1 summarizes the analyses of the EEG data. The analyses included measures of oscillatory brain activity (power) and connectivity in electrode and source space, respectively. Two categories of analyses were performed. First, local, i.e. spatially specific, analyses were performed in which a single value is obtained for every electrode, voxel or brain region. These analyses include the comparison of power topographies on electrode level and connectivity and local network measures topographies (degree, clustering coefficient) on source level. Second, global, i.e. spatially holistic, analyses were performed. These analyses include all analyses which average across all electrodes, voxels or brain regions, i.e. the peak frequency, the power spectrum and all global network measures (see below). In all global analyses, each participant is represented by a single scalar value per measure. All analyses were based on 2-second epochs of resting-state data to balance robustness, frequency resolution and non-stationarity of the data.

**Figure 1.**
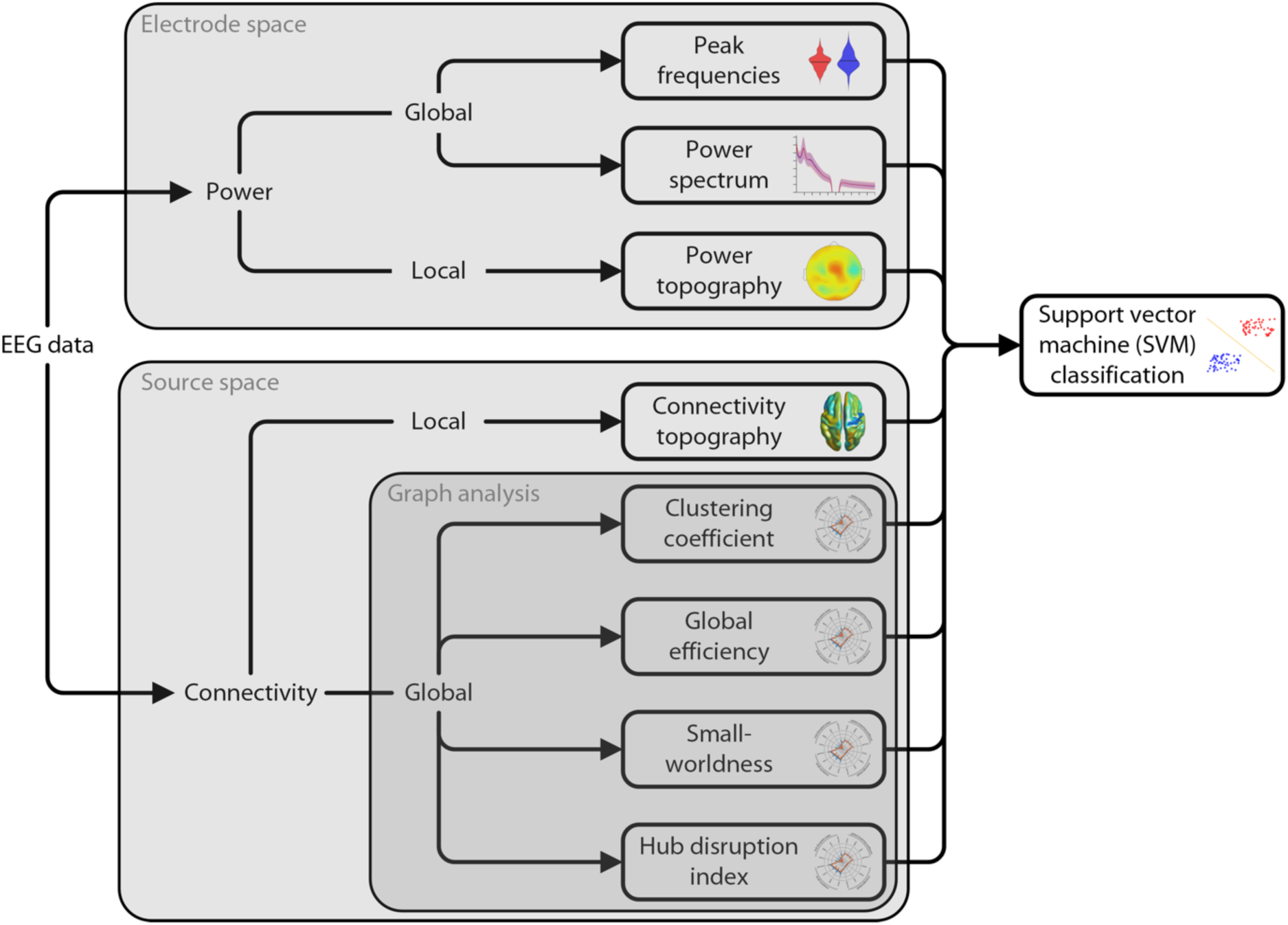
Analysis pipeline. EEG data were analyzed with regards to power and connectivity, which quantify neural activity and neural communication, respectively. Power analyses were performed in electrode space. Analyses of functional connectivity were performed in source space. Connectivity analyses comprised phase-based (PLV, dwPLI) and amplitude-based (AEC) connectivity measures. Graph theoretical network analysis was applied to further characterize functional connectivity. All measures were compared between chronic pain patients and healthy controls. In addition, a purely data-driven machine learning approach was adopted, using SVMs. The SVM was trained on all power and connectivity measures to distinguish between chronic pain patients and healthy controls.

### Brain activity (power) analysis

Oscillatory brain activity (power) was analyzed in electrode space. Frequency-specific global power was computed for all epochs using a Fast Fourier Transformation with Slepian multitapers and a frequency smoothing of 1 Hz and then averaged across epochs and electrodes. Power was first computed for the complete power spectrum, i.e. 1 – 100 Hz, with a frequency resolution of 0.5 Hz. To remove line noise, a band-stop filter of 45 – 55 Hz was applied before computing power.

The individual dominant peak frequency was determined on the average of the epochs as the highest local maximum (larger than its two neighboring samples) of the amplitude in the frequency range of 6 – 14 Hz [7]. We also pursued alternative approaches to determine the peak frequency by (i) computing the center of gravity of the power spectrum [7; 41], (ii) computing the dominant peak frequency using longer time windows of 5 s, and by (iii) computing the peak frequency on each single epoch and then averaging the peaks [31].

To compare the spatial distribution of local brain activity between patients and healthy controls, power was averaged within each frequency band before comparing frequency-specific power topographies between groups using cluster-based permutation tests.

Relative power was calculated by normalizing every power value (both local and global power) by the respective participant’s total power. Total power was calculated by summing all power values across frequencies from 1 to 100 Hz and across all electrodes.

### Connectivity analysis

Connectivity analyses were performed in source space. For source analysis, we used linearly constrained minimum variance beamforming [82] to project the band-pass filtered data for each frequency band and participant from electrode space into source space. Spatial filters were computed based on the covariance matrices of the band-pass filtered data for each frequency band and a lead field matrix. A three-dimensional grid with a 1 cm resolution covering the brain was defined, resulting in a total of 2020 voxels in the brain. The lead field was constructed for each voxel using a realistically shaped three-shell boundary-element volume conduction model based on the template Montreal Neurological Institute (MNI) brain. We used a regularization parameter of 5 percent of the covariance matrix and chose the dipole orientation of most variance using singular value decomposition. Finally, the preprocessed and bandpass-filtered EEG data were projected through the spatial filter to extract the amplitude time series of neuronal activity of each frequency band at each voxel.

Connectivity analyses of EEG data were performed using phase-based and amplitude-based approaches which capture different and complementary neural communication processes [23]. Amplitude-based connectivity is more closely related to structural connectivity [86; 91] and putatively regulates activation of brain regions [23]. Phase-based connectivity seems more detached from structure and is more strongly affected by contextual factors [70; 71]. Here, functional connectivity was investigated using the phase locking value (PLV) [45], the debiased weighted phase lag index (dwPLI) [87] and the orthogonalized amplitude envelope correlation (AEC) [36]. The PLV and dwPLI are based on the phase of the signals, whereas the AEC is based on the amplitude. The PLV is well-established, highly sensitive and captures both zero phase lag and non-zero phase lag connectivity. The dwPLI captures non-zero phase lag connectivity only. The dwPLI is therefore not susceptible to volume conduction at the cost of reduced sensitivity because real synchrony at zero phase lag is also discarded. Because of the explorative character of our study, we mainly report the more sensitive measure, the PLV. All our connectivity analyses are based on contrasts between two groups and therefore less susceptible to effects of volume conduction.

For the connectivity analyses, the connectivity between every pair of voxels was computed, resulting in a 2020 × 2020 matrix, with a single value representing the strength of connection between two voxels over the complete recording time. All three connectivity measures are undirected.

### Graph-theoretical network analysis

By applying graph-theoretical methods to the data, we reduced the high-dimensional EEG data to a few network measures, characterizing the organization of the whole brain network. Graph theory defines networks as collections of *nodes* and *edges* connecting the *nodes* to each other. We defined the *nodes* as voxels and the *edges* as thresholded functional connectivity between voxels. To reduce the computational load, the adjacency matrix (the matrix defining all *edges* between the *nodes*) was thresholded to the 10% (5%, 20%) strongest connections and binarized, resulting in an unweighted and undirected graph.

We used common graph measures that characterize either a single node (local analyses) or the complete network (global analyses) [64]. We investigated two local network measures: the *degree* and the *local clustering coefficient*. The *degree* is the number of a node’s edges, i.e. the number of connections to other nodes. The *local clustering coefficient* (CC) is the fraction of the node’s neighbors that are also neighbors of each other. Thus, both local measures depict the importance of a node. Four global network measures were included in the analysis: the *global clustering coefficient* (gCC), *global efficiency* (gEff), *small-worldness* (S) and *hub disruption index* (k_d_) [2]. The *global clustering coefficient* is the average of the local clustering coefficient of all nodes, resulting in a measure of local segregation, i.e. how specialized sub-regions of the brain are [28; 64]. The *global efficiency* is the inverse of the average shortest path length. It measures the strength of “long-range” connections or the global integration [1; 3; 46; 64]. *Small-worldness* describes the ratio of *clustering coefficient* and *global efficiency* and compares it to random networks. This measure has been associated with efficiency of communication in a network [92]. Lastly, the *hub disruption index* compares the *degree* of all nodes to those of a control group. Positive values indicate that strongly connected nodes increase and weakly connected nodes decrease their number of connections (“the rich get richer and the poor get poorer”). Conversely, negative values indicate that strongly connected nodes decrease and weakly connected nodes increase their number of connections (“the rich get poorer and the poor get richer”), which means a shift of the network towards a random network with less internal structure [2; 50; 51].

### Correlation analysis

Pearson’s r was computed between clinical parameters and brain measures, which were found to show significant relationships either in previous studies [13; 20; 24; 33; 44; 51; 65; 66; 76; 83] or our own. The global peak frequency, mean global power in the four frequency bands, hub disruption index and the mean theta and gamma PLV connectivity, the PLV global efficiency in the gamma band and the dwPLI hub disruption index in the gamma band, were correlated with the following clinical parameters: current pain intensity, average pain intensity in the last four weeks quantified by a visual analog scale, pain duration, pain disability quantified by the PDI [21], mental and physical quality of life quantified by the VR-12 [68], depression quantified by the BDI-II [8] and medication as quantified by the medication quantification scale (MQS) [34].

### Machine learning analysis

The machine learning analysis was carried out using the Statistics and Machine Learning Toolbox in Matlab as well as custom-written scripts. We implemented a support vector machine (SVM) [16] with a linear kernel, which was trained on all available features. The features were the peak frequency (one feature per participant), global power spectrum (199 features per participant, one feature for each frequency step), local power (4 × 65 features per participant, 65 electrodes), local strength of connectivity (3 × 4 × 2020 features per participant, 2020 voxels in 4 frequency bands and 3 connectivity measures), local degree (3 × 4 × 2020 features per participant, 2020 voxels in 4 frequency bands and 3 connectivity measures), local clustering coefficient (3 × 4 × 2020 features per participant, 2020 voxels in 4 frequency bands and 3 connectivity measures), and the global graph measures (3 × 4 × 4 features per participant, 4 global graph measures in 4 frequency bands and 3 connectivity measures). This resulted in an SVM containing 73228 features per participant. To avoid overfitting, we implemented a so-called sequential forward feature selection. This algorithm randomly adds single features to the SVM and evaluates the performance. When performance is increased, the feature is retained, and another feature is added. If performance of the SVM is not improved, the feature is discarded, and a different feature is added. This process is repeated until no additional feature improves performance. The performance of the SVM was evaluated using a 10-fold cross-validation. First, the dataset was randomly split into 10 folds. 9/10 folds were used to train the classifier, with a nested feature selection loop, after which the remaining 1/10 were classified, resulting in a certain classification accuracy. This procedure was then repeated, cycling through all folds, yielding a mean accuracy over the 10 folds. As our groups were unbalanced regarding participant numbers, we randomly picked 84 datasets from the cohort of chronic pain patients for the classification procedure, repeating this 1000 times. To conclude whether this result truly exceeds chance level, we repeated the whole procedure with the same data, but the labels of chronic pain patients and healthy controls were randomly shuffled [15]. The two resulting distributions were then statistically compared using a non-parametric permutation test. The sensitivity is defined as the rate of true positives, i.e. correctly classified patients, divided by the number of total patient classifications. The specificity is defined as the rate of true negatives, i.e. correctly classified healthy controls, divided by the number of total healthy classifications.

Apart from the overall performance of the SVM, we also investigated which features contained the highest predictive value, i.e. which features were most consistently picked by the SVMs by looking at the number of times a certain feature was included in the SVM by automatic feature selection.

### Statistical analysis

Statistical analysis was carried out using FieldTrip [57] and custom-written Matlab scripts. The significance level for all statistical tests was set to 0.05 two-tailed. We used cluster-based permutation tests with a cluster-threshold of 0.05 to compare patients to healthy controls for all local analyses and the global power spectrum analysis [52]. The other global measures were compared using non-parametric permutation tests, permuting between the patient and healthy control group. The underlying statistical test for all permutation tests was an unpaired T-test. To account for multiple comparisons within a specific measure, we corrected all p-values of tests that were performed on all 4 frequency bands using the Holm-Bonferroni method [37]. To test whether the accuracy of the SVMs was above random chance level, we computed a non-parametric permutation test on the accuracy distribution of the SVM trained on the real data against the accuracy distribution of the SVM trained on the randomly labeled data. Pearson’s r was calculated to find correlations between brain measures and clinical parameters and tested for statistical significance against the null hypothesis of no correlation. Resulting p-values were again corrected for multiple comparisons across the four frequency bands using the Holm-Bonferroni method, if applicable.

### Data availability

All scripts and data are available upon request.

## Results

### Global measures of brain activity

We first investigated whether chronic pain was associated with global changes of oscillatory brain activity. We determined the peak frequency of EEG activity in chronic pain patients and healthy controls by averaging the power spectra across all epochs and electrodes and determining the maximal power in the frequency range of 6 to 14 Hz. Peak frequency was 9.8 ± 1.2 Hz (mean ± standard deviation) in chronic pain patients and 10.0 ± 1.4 Hz in healthy controls and did not differ significantly between groups (non-parametric permutation test, p = 0.20, Fig. 2A). Other common approaches to determine the peak frequency, by computing the Center of Gravity, analyzing longer time windows (5 seconds) for increased spectral resolution, or computing the peak frequency on individual epochs and then averaging, did not show a difference between groups either (non-parametric permutation tests, p ≥ 0.14).

**Figure 2.**
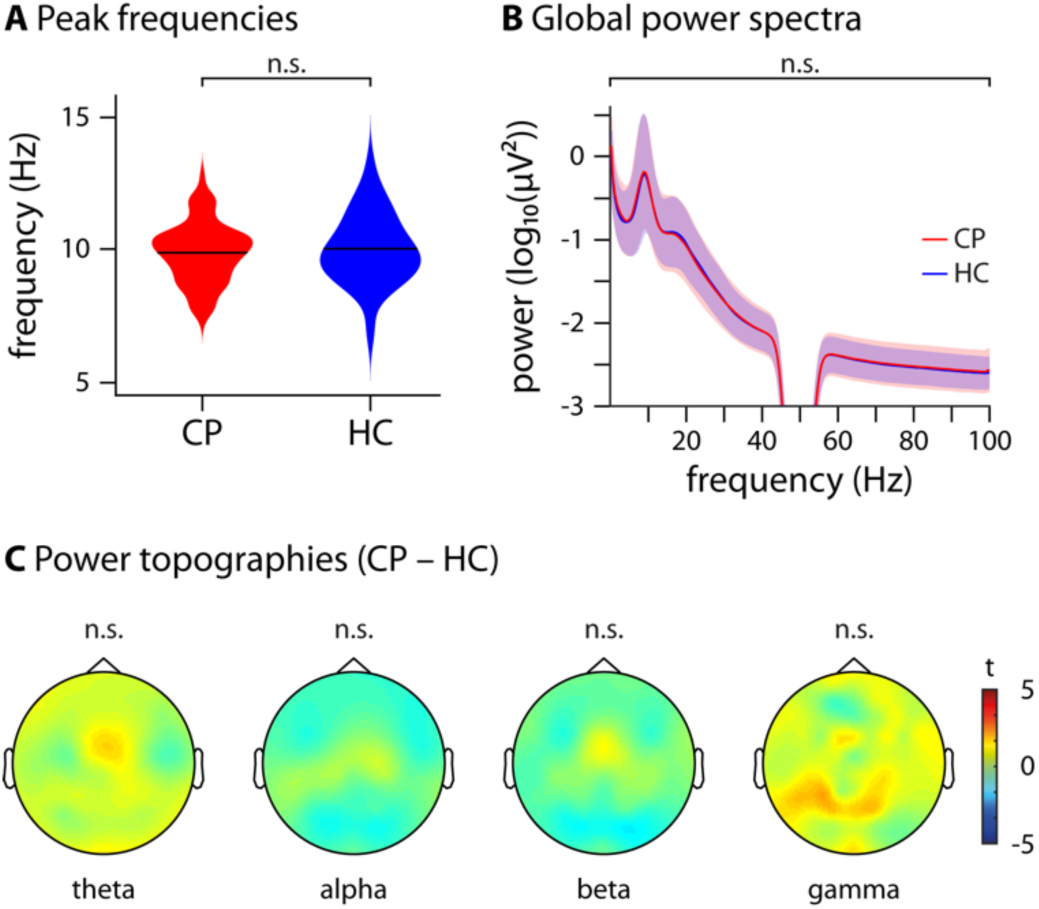
Global and local measures of brain activity. **(A)** Violin plot of the dominant peak frequencies computed on the average across all electrodes of chronic pain patients (CP, red) and healthy controls (HC, blue). A non-parametric permutation test showed no significant difference (p = 0.20) between the two groups. **(B)** Global power spectra of CP (red) and HC (blue), averaged across all electrodes and shown for the frequencies 1 – 100 Hz, with a bandstop filter at 45 – 55 Hz to remove line noise. A cluster-based permutation test clustered across frequencies did not show any significant differences (t_max/min = 1.7/−1.5). **(C)** Scalp topographies of power differences between CP and HC at theta, alpha, beta and gamma frequencies, averaged across frequencies in each band. The colormap shows the t-values of a cluster-based permutation test. No significant clusters were found in any frequency band (theta t_max/min = 1.5/−0.6, alpha t_max/min = 0.62/−1.2, beta t_max/min = 1.0/−1.4, gamma t_max/min = 2.1/−0.7).

Next, we examined whether chronic pain was associated with global changes of oscillatory brain activity at any frequency between 1 and 100 Hz. To this end, we compared the global power spectrum of EEG activity averaged across all electrodes between chronic pain patients and healthy controls. The results did not reveal any significant difference between the two groups at any frequency (cluster-based permutation statistics clustered across frequencies, t_max/min = 1.7/−1.5; Fig. 2B). When controlling for inter-subject differences in overall power by calculating power relative to the total power across all electrodes and frequencies for each participant, the results did not show a significant difference between patients and controls either (t_max/min = 1.4/−1.7).

Thus, we did not observe global changes of the peak frequency or the power spectrum of oscillatory brain activity in chronic pain patients.

### Local measures of brain activity

We further assessed whether chronic pain was associated with local changes of oscillatory brain activity. We therefore calculated topographical maps of brain activity for theta (4 – 8 Hz), alpha (8 – 13 Hz), beta (14 – 30 Hz) and gamma (60 – 100 Hz) frequency bands. Group comparisons of the topographical maps did not show significant differences between patients and controls in any frequency band (cluster-based permutation statistics clustered across electrodes, t_max/min = 2.0/−1.4, Fig. 2C). When controlling for inter-subject differences in overall power by calculating relative power, the results did not show a significant difference between patients and controls either (t_max/min = 2.5/−3.2).

Thus, our findings did not show local changes of oscillatory brain activity in chronic pain patients in any frequency band.

### Local measures of functional connectivity

Next, we explored whether chronic pain was associated with changes of functional connectivity which is a measure of neural communication. To reduce potential confounds by field spread and volume conduction effects [67], we performed all connectivity analyses in source space using 2020 voxels with a size of 1 × 1 × 1 cm^3^. We first investigated whether chronic pain was associated with local changes of functional connectivity in any brain region or any frequency band. To this end, we calculated the *connectivity strength* for each voxel and frequency band. *Connectivity strength* was defined as the average connectivity of a specific voxel to all other voxels of the brain, which yields one *connectivity strength* value for each voxel. This allows for visualizing *connectivity strength* in a topographical map and applying the same statistical approaches used for the analysis of local brain activity. Analysis of phase-based connectivity (Fig. 3A) showed that chronic pain patients’ *connectivity strength* in the theta band was significantly increased (cluster-based permutation test, p(corrected/uncorrected) = 0.045/0.011, Cohen’s d = 0.40) in comparison to the control group. The strongest contrast was found in the supplementary motor area (MNI = [−10, 10, 70]). Moreover, we also found that patients showed a significantly increased *connectivity strength* in the gamma band (cluster-based permutation test, p(corrected/uncorrected) = 0.0072/0.0018, Cohen’s d = 0.59), which was maximal in the anterior prefrontal cortex (MNI = [−40, 40, 30]). As in both frequency bands only a single cluster of increased connectivity was found, the increase likely reflects local frontal connectivity within the clusters rather than connectivity to targets outside of the clusters. No significant clusters were found in the alpha and beta bands (alpha: p_min(corrected/uncorrected) = 1/0.61, t_max = 2.8; beta: p_min(corrected/uncorrected) = 0.71/0.18, t_max = 3.5). Analysis of amplitude-based connectivity did not show any significant differences in *connectivity strength* between chronic pain patients and healthy controls in any brain region or any frequency band. (Fig. 3B, t_max/min = 0.4/−1.2).

**Figure 3.**
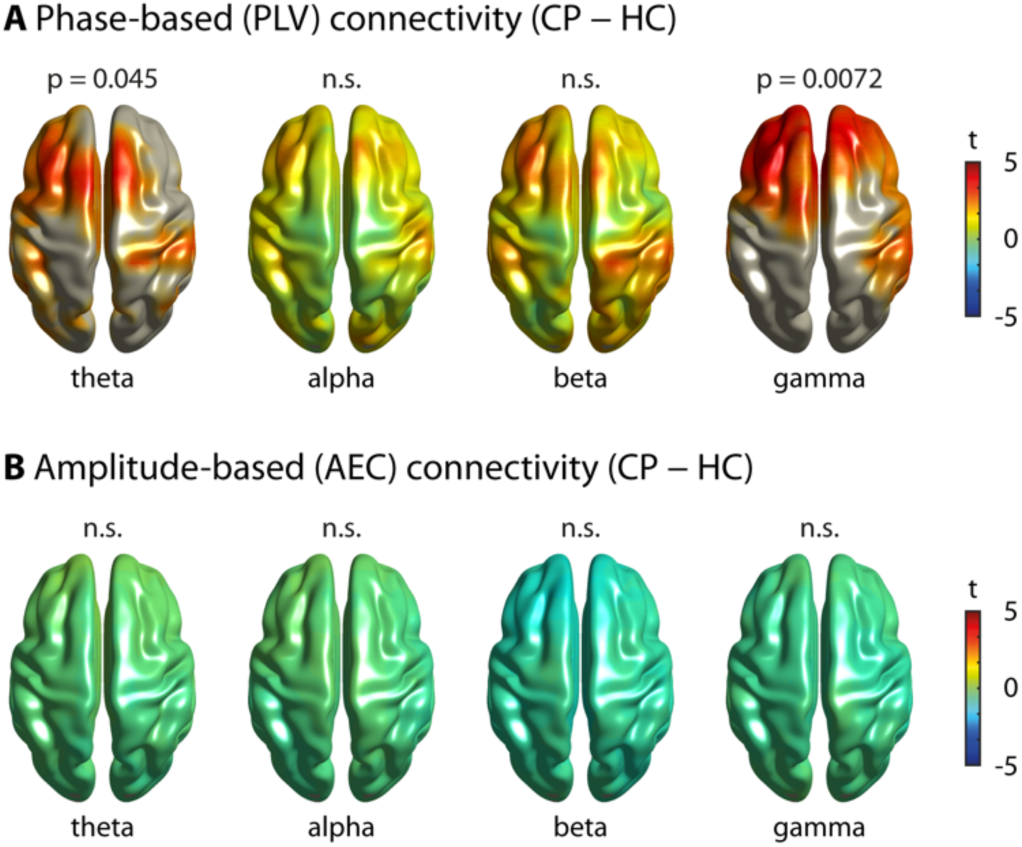
Local measures of functional connectivity. Brain topographies of the comparison of connectivity strength between chronic pain patients (CP) and healthy controls (HC) in the theta, alpha, beta and gamma band frequencies, averaged across frequencies in each band, are shown. Connectivity strength was calculated as the average connectivity of one voxel to all other voxels of the brain. The colormaps show the t-values. Significant results are masked, i.e. all voxels but the ones belonging to a significant cluster are greyed out. When no significant clusters are found, nothing is greyed out to show potential trends. **(A)** Phase-based connectivity (phase locking value, PLV). A significant increase of chronic pain patients’ connectivity strength was observed in the theta band (p(corrected/uncorrected) = 0.045/0.011, t_max = 3.8, Cohen’s d = 0.40) and the gamma band (p(corrected/uncorrected) = 0.0072/0.0018, t_max = 5.1, Cohen’s d = 0.59). **(B)** Amplitude-based connectivity (orthogonalized amplitude envelope correlation, AEC). No significant differences were found in any frequency band (theta t_max/min = 0.4/−0.6, alpha t_max/min = 0.1/−0.7, beta t_max/min = −0.3/−1.2, gamma t_max/min = 0.0/−1.1).

To further assess connectivity patterns of chronic pain patients, we performed graph theory-based network analysis. We first examined the local properties of brain networks in chronic pain patients. A basic property of a node is the number of its connections to other nodes, which is termed the *degree*. Conceptually, the *degree* is closely related to the *connectivity strength* analyzed in the previous step, the essential difference being that the edges are thresholded, whereas the *connectivity strength* considers all connections. We computed the *degree* of every voxel and compared it between patients and controls. No difference in *degree* was found in any frequency band. This applied similarly to phase-based and amplitude-based measures of connectivity (PLV: p_min(corrected/uncorrected) = 0.51/0.13, t_max = 4.2; AEC: p_min(corrected/uncorrected) = 1/0.56, t_max = 3.0). This lack of a difference in *degree* indicates that the difference in *connectivity strength* is not confined to the strongest connections but instead applies to connections of all strengths.

Another well-established measure that characterizes nodes is the *clustering coefficient*. This measure assesses the number of connections of neighboring nodes, i.e. it measures the local integration of a node served by short-range connectivity. Comparing the *clustering coefficients* of all nodes between patients and controls did not show any significant differences at any frequency band, neither for phase-based nor amplitude-based connectivity (PLV: p_min(corrected/uncorrected) = 0.12/0.030, t_max = 3.3, AEC: p_min(corrected/uncorrected) = 1/0.28, t_max = 3.8).

Taken together, the analysis of local measures of functional connectivity showed increases of phase-based connectivity in frontal and prefrontal cortices at theta and gamma frequencies in chronic pain patients. The increase in the theta band was of small effect size (Cohen’s d = 0.40), whereas the increase in the gamma band was of medium effect size (Cohen’s d = 0.59).

### Global measures of functional connectivity

We next investigated whether chronic pain was associated with changes of global measures of functional connectivity and therefore computed graph measures which characterize different and complementary global properties of brain networks. Figure 4 summarizes the results of the global graph measures. First, we calculated the *global clustering coefficient*, which is commonly interpreted as a measure of functional segregation in a network. We found no differences in *global clustering coefficient* between chronic pain patients and healthy controls (Table 3; p_min (corrected/uncorrected) = 0.088/0.022). Second, we assessed the *global efficiency*, which provides an account of the ease of long-distance communication and is commonly interpreted as a measure of functional integration in a network. After accounting for multiple comparisons, we found evidence for a decrease of *global efficiency* in patients in the gamma frequency band when investigating phase-based connectivity (Table 3; p(corrected/uncorrected) = 0.013/0.0032). The effect size of this difference was small (Cohen’s d = 0.44). Third, we computed the *small-worldness*, which is associated with communication efficiency within a network. We detected no changes in *small-worldness* between the two groups (Table 3; p_min(corrected/uncorrected) = 0.26/0.064). Fourth, we analyzed the *hub disruption index*, which has been shown to differ between chronic pain patients and controls in previous functional magnetic resonance imaging studies [50; 51]. It shows potential shifts of connections which manifest on a global scale. Our results did not show a difference of the *hub disruption index* in any frequency band when comparing chronic pain patients to healthy controls (Table 3; p_min(corrected/uncorrected) = 1/0.32).

**Figure 4.**
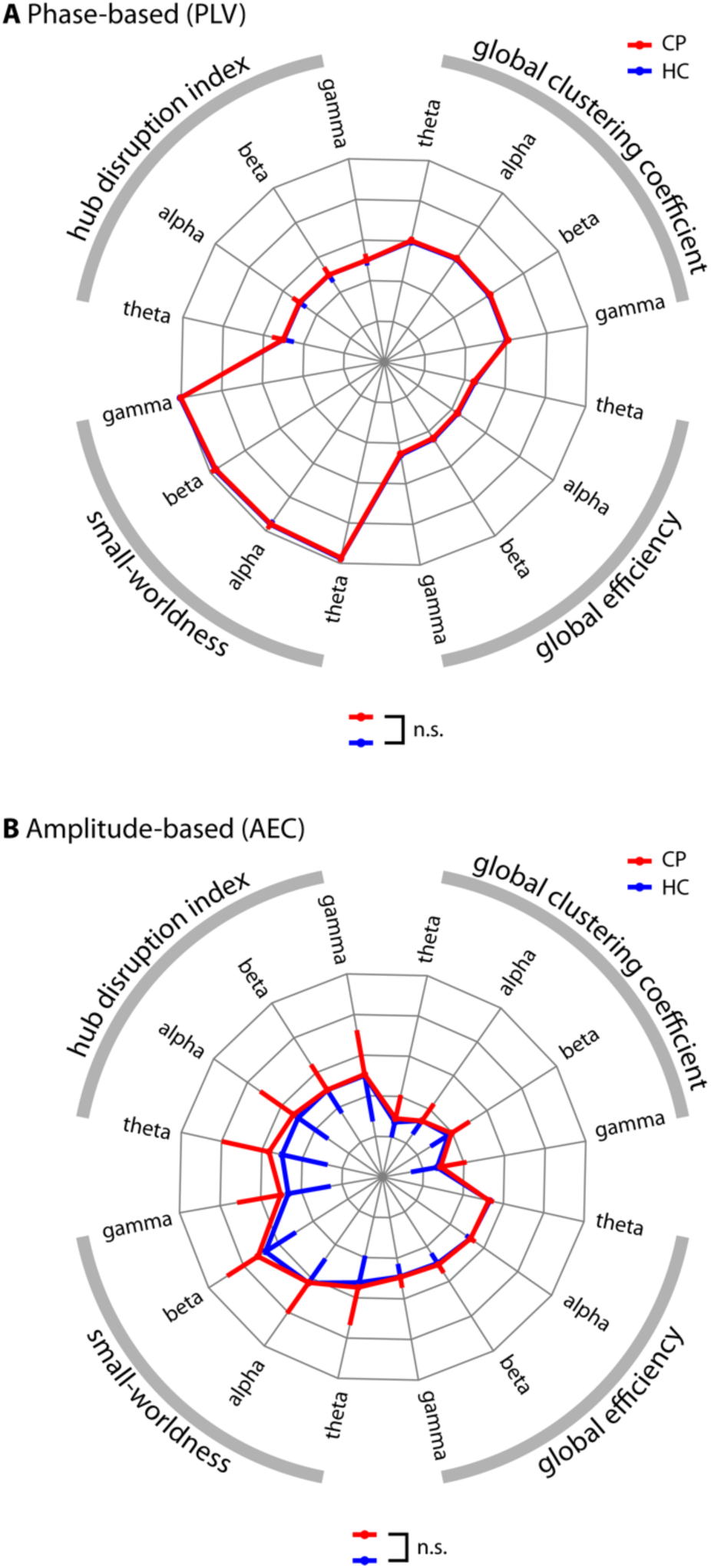
Global graph theoretical measures of functional connectivity. The radar plots show four global graph measures in four frequency bands based on **(A)** phase-based and **(B)** amplitude-based connectivity measures. The clockwise arrangement follows the following pattern: theta, alpha, beta and gamma repeated for the four graph measures: global clustering coefficient, global efficiency, small-worldness and absolute values of the hub disruption index. The red lines show the chronic pain patients’ (CP) values, while the blue lines represent the healthy controls’ (HC). Error bars show the standard deviation. For visualization purposes, the symmetric error bars are only drawn in a single radial direction. Axes run from the center (= 0) to the outside (= 1). For visualization purposes, the small-worldness and hub disruption index were scaled with a factor of 0.2. **(A)** Phase-based connectivity (phase locking value, PLV). The global efficiency in the gamma band was significantly decreased in chronic pain patients (non-parametric permutation test, p(corrected/uncorrected) = 0.013/0.0032, Cohen’s d = 0. 44). No other measure revealed a significant difference when compared between groups, see Table 3 for details. **(B)** Amplitude-based connectivity (orthogonalized amplitude envelope correlation, AEC). No significant difference between groups was observed, see Table 3 for statistical details.

**Table 3.** Comparisons of global graph measures of functional connectivity between chronic pain patients and healthy controls. Uncorrected p-values of non-parametric permutation tests comparing global graph measures between groups. P-values were corrected for multiple comparisons using the Holm-Bonferroni method across the four frequency bands to take cross-spectral dependencies into account. After correction, only the PLV-based global efficiency in the gamma band differed significantly between groups (p(corrected) = 0.013, Cohen’s d = 0.44). Color of cells indicate the direction of significant effects, blue indicates lower values in chronic pain patients. PLV, phase locking value; AEC, orthogonalized amplitude envelope correlation; gCC, global clustering coefficient; gEff, global efficiency; S, small-worldness; k_d_, hub disruption index.

|  | PLV |  |  |  | AEC |  |  |  |
| --- | --- | --- | --- | --- | --- | --- | --- | --- |
|  | theta | alpha | beta | gamma | theta | alpha | beta | gamma |
| gCC | 0.084 | 0.040 | 0.092 | 0.022 | 0.22 | 0.92 | 0.29 | 0.38 |
| gEff | 0.052 | 0.13 | 0.084 | 0.0032 | 0.17 | 0.78 | 0.30 | 0.74 |
| S | 0.17 | 0.24 | 0.24 | 0.064 | 0.30 | 0.92 | 0.094 | 0.24 |
| k<sub>d</sub> | 0.77 | 0.64 | 0.94 | 0.88 | 0.066 | 0.32 | 0.99 | 0.88 |

In summary, global graph measures of functional connectivity showed a decrease of *global efficiency* at gamma frequencies in chronic pain patients. This decrease was of small effect size (Cohen’s d = 0.44).

### Additional functional connectivity analyses

First, we tested whether changes of functional connectivity in chronic pain patients can be detected using another common phase-based connectivity measure (debiased weighted phase lag index, dwPLI)[75; 87]. The dwPLI differs from the PLV by capturing non-zero phase lag connectivity only. The dwPLI is therefore not susceptible to volume conduction which can yield artificial connectivity effects at the cost of reduced sensitivity because real synchrony at zero phase lag is also discarded. The results of the cluster-based permutation tests did not reveal any local difference in functional connectivity between patients and controls (p_min (corrected/uncorrected) = 0.48/0.12, t_min = −3.0). This indicates that zero phase lag connectivity plays an important role in the increased frontal connectivity of patients. Concerning global graph measures (Table 4), the *hub disruption index* was significantly lower in chronic pain patients in the gamma band (p(corrected/uncorrected) = 0.00/0.00, Cohen’s d = 0.63).

**Table 4.** Comparisons of global graph measures between chronic pain patients and healthy controls - debiased weighted phase lag index (dwPLI). Uncorrected p-values of non-parametric permutation tests comparing global graph measures between patients with chronic pain and healthy controls. P-values were corrected for multiple comparisons using the Holm-Bonferroni method across the four frequency bands to take cross-spectral dependencies into account. After correction, only the dwPLI-based hub disruption index (k_d_) in the gamma band was significantly lower in patients (p(corrected) = 0.00, Cohen’s d = 0.63). Color of cells indicate the direction of significant effects, blue indicates lower values in chronic pain patients. gCC, global clustering coefficient; gEff, global efficiency; S, small-worldness; k_d_, hub disruption index.

|  | dwPLI |  |  |  |
| --- | --- | --- | --- | --- |
|  | theta | alpha | beta | gamma |
| gCC | 0.088 | 0.45 | 0.017 | 0.47 |
| gEff | 0.29 | 0.051 | 0.20 | 0.12 |
| S | 0.085 | 0.30 | 0.090 | 0.29 |
| k<sub>d</sub> | 0.048 | 0.20 | 0.43 | 0.00 |

Second, we performed control network analyses by calculating all graph measures with different edge densities using all three connectivity measures. This was done in order to examine the robustness of our results, which were based on a threshold of 10% strongest connections. No significant group differences in local graph measures were found for any of the connectivity measures. Regarding global measures, the *global efficiency* in the gamma band calculated with the PLV (5% edge density, p(corrected) = 0.011, Cohen’s d = 0.44; 20% edge density, p(corrected) = 0.0080, Cohen’s d = 0.44) and the hub disruption index in the gamma band calculated with the dwPLI (5% edge density, p(corrected) = 0.00, Cohen’s d = 0.61; 20% edge density, p(corrected) = 0.00, Cohen’s d = 0.61) were both significantly lower in chronic pain patients irrespective of edge density and therefore showed a consistent and robust effect.

Third, we tested whether depression plays a critical role in explaining differences between patients and controls. We therefore aimed to replicate the significant findings after excluding patients with a clinically relevant depression (BDI-II score ≥ 18, n = 36). This re-analysis did not qualitatively change any of the previously significant results. Frontal connectivity was increased for chronic pain patients without depression in the theta (p(corrected) = 0.0080, Cohen’s d = 0.65) and gamma (p(corrected) = 0.0048), Cohen’s d = 0.68) bands. Similarly, PLV-based global efficiency (p(corrected) = 0.014, Cohen’s d = 0.42) and the dwPLI-based hub disruption index (p(corrected) = 0.00, d = 0.75) were decreased at gamma frequencies in chronic pain patients without depression.

In summary, the PLV *global efficiency* and the dwPLI *hub disruption index* in the gamma band were consistently changed in chronic pain patients even when excluding patients with depression. Both measures were decreased in chronic pain patients, the PLV global efficiency showing a small effect size (average Cohen’s d = 0.44) and the dwPLI hub disruption index showing a medium effect size (average Cohen’s d = 0.61).

### Relationship between brain activity/functional connectivity and clinical parameters

We further investigated the relationships of brain-based activity and connectivity measures with clinical parameters. To reduce the number of statistical tests, we restrained our analyses to selected measures of brain activity and brain connectivity, which were associated with clinical parameters of chronic pain patients in previous studies [13; 20; 24; 33; 44; 51; 65; 66; 76; 83]. We thus computed correlations between the global peak frequency, mean global power in the four frequency bands, the *hub disruption index* and the following major clinical parameters: current pain intensity, average pain intensity in the last four weeks, pain duration, pain disability, mental and physical quality of life, depression and medication as quantified by the medication quantification scale. Additionally, we correlated the significant clusters in the theta and gamma PLV connectivity, the PLV *global efficiency* in the gamma band and the dwPLI *hub disruption index* with the same clinical parameters. The results showed no significant correlations (Fig. 5). Thus, we did not observe any relations of measures of brain activity and functional connectivity with clinical parameters including medication. This suggests that increases of frontal connectivity and global network changes in chronic pain patients do not scale with disease characteristics but rather characterize the state of chronic pain *per se*.

**Figure 5.**
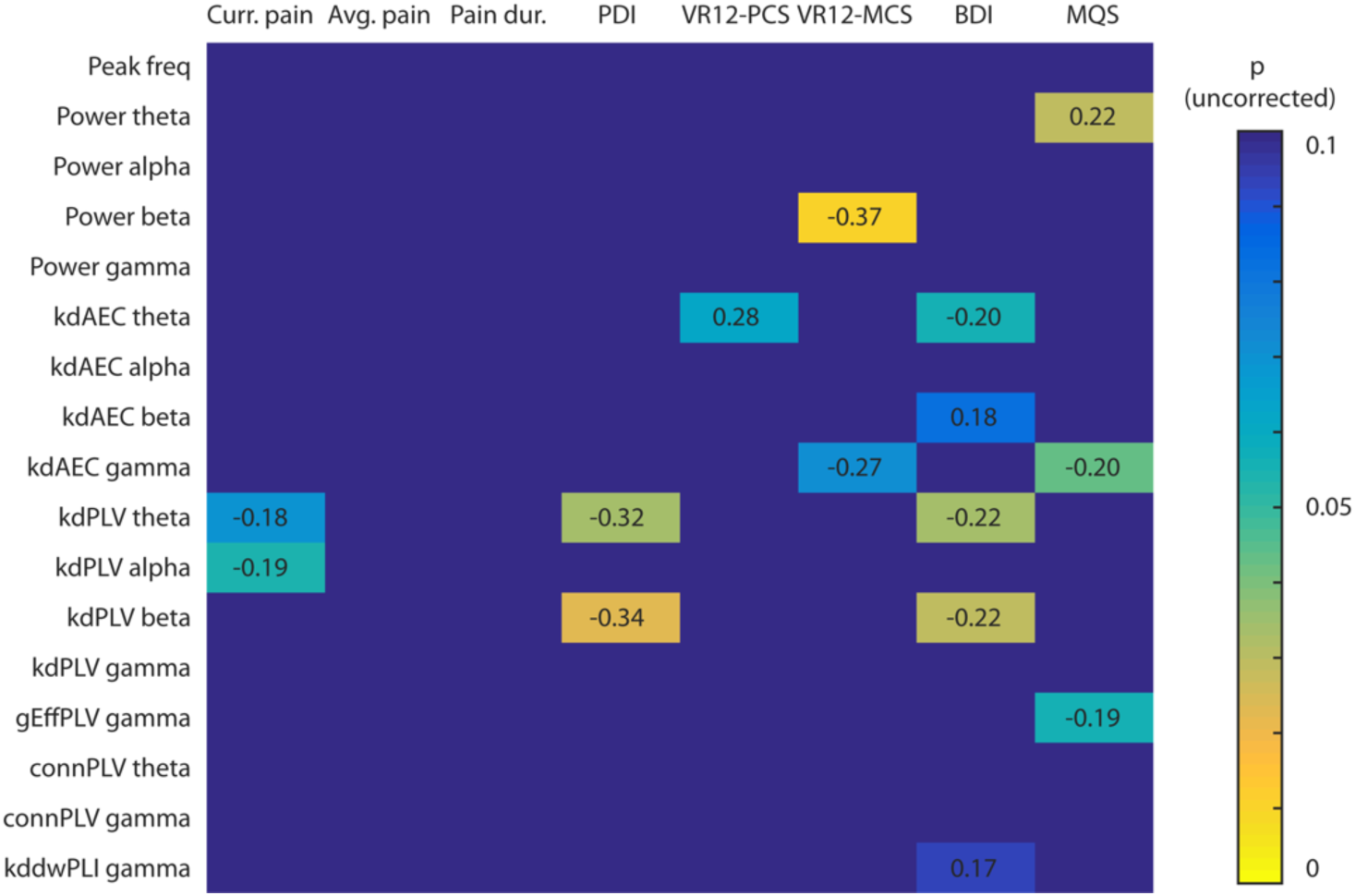
Correlations between clinical/behavioral parameters and brain activity/functional connectivity measures. The cell values show the strength and direction of the correlations (Pearson’s r) and the color depicts the uncorrected p values. Only correlations showing a trend (p < 0.1) are shown. No correlation was statistically significant after Holm-Bonferroni correction for multiple comparisons across the four frequency bands. Curr. pain, current pain intensity; Avg. pain, average pain intensity in the last 4 weeks; Pain dur., pain duration; PDI, pain disability index; VR12-PCS, Veterans’s RAND physical component score; VR12-MCS, Veterans’s RAND mental component score; BDI, Beck Depression Inventory II, MQS, medication quantification scale; peak freq, peak frequency; k_d_, hub disruption index; AEC, measure is based on the orthogonalized amplitude envelope correlation; PLV, measure is based on the phase locking value; dwPLI, measure is based on the debiased weighted phase lag index; gEff, global efficiency; conn, connectivity strength.

### Machine learning approach

Finally, we performed a multivariate machine learning approach. This approach extends the previous univariate approaches by taking patterns of brain activity and connectivity into account rather than single pieces of information in isolation. Moreover, it complements the previous descriptive group analyses by adding a predictive, single-subject analysis. We used an SVM classifier to test whether patterns of brain activity and/or connectivity can distinguish between chronic pain patients and healthy controls. We trained a linear SVM on all aforementioned measures of brain activity and functional connectivity, using an automated sequential feature selection algorithm. The performance of the SVM was evaluated using a 10-fold cross-validation. The resulting mean accuracy was 57% ± 4% with a sensitivity of 60% ± 5% and a specificity of 57% ± 5%. To test whether this result exceeds chance level, we repeated the whole procedure with the same data but randomly shuffled labels of chronic pain patients and healthy controls. This resulted in a permutation distribution with 50% ± 5% accuracy. A non-parametric permutation test of the 2 accuracy distributions (Fig. 6A) confirmed that the real model was significantly more accurate than random guessing (p < 0.001). Finally, we were interested to know which features of brain activity and/or connectivity were most relevant for the classification. The automatic feature selection on average picked 5.5 features for the SVMs. We therefore show the five most frequently picked features in Figure 6B. The most relevant features were phase-based connectivity measures in frontal brain areas at gamma (MNI: −40, 30, 40 and −30, 50, 10) and theta (MNI: −20, 50, 40) frequencies. These features were chosen with a rate of about 15% each. Thus, in more than 50 % of classifications, phase-based connectivity in frontal brain areas was the most relevant feature.

**Figure 6.**
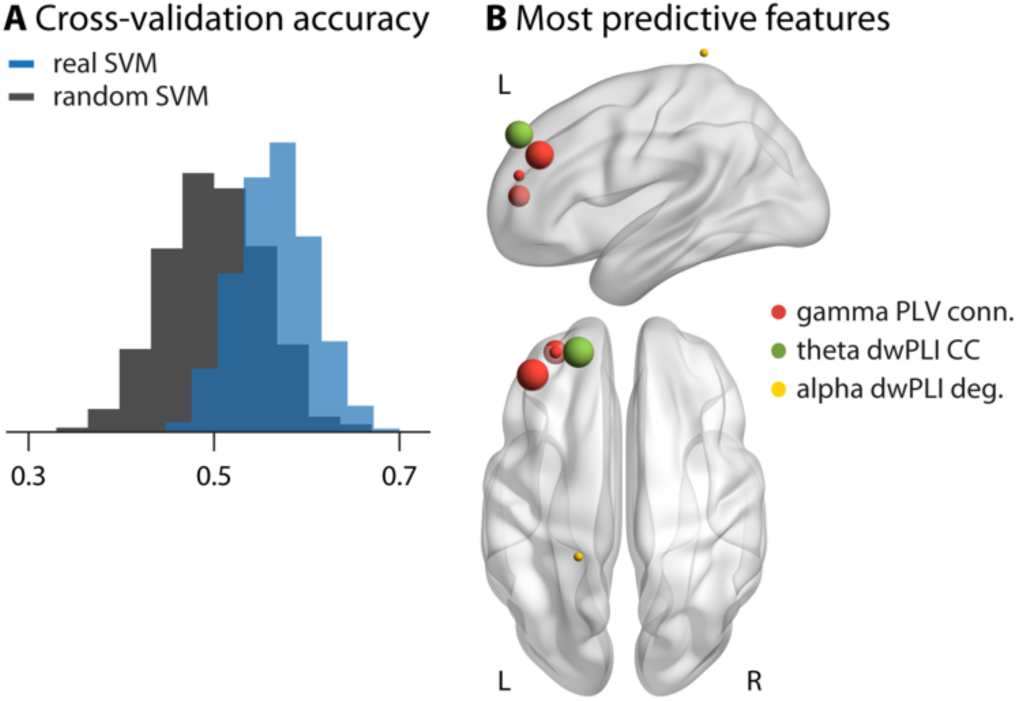
Multivariate machine learning approach to classify chronic pain patients and healthy controls. **(A)** Distribution of mean accuracies resulting from a 10-fold cross-validation. The blue histogram shows the results trained on the actual data including all features of brain activity and connectivity. The gray histogram shows a support vector machine (SVM) trained on data with randomly permuted labels. The SVM trained on the real data shows an accuracy of 57 ± 4 %, significantly higher than the accuracy of the SVM trained on randomly permuted data, 50 ± 5 % (p < 0.001). **(B)** The 5 most predictive features, i.e. those selected most consistently by the SVMs. Specific measures are color-coded, the size of the spheres represents how often a specific feature was selected. The most frequently selected features were phase locking value (PLV) based connectivity of the prefrontal cortex (MNI: −40, 30, 40 and −30, 50, 10) in the gamma band, which were selected in 15 % and 12 % of SVMs, respectively, and debiased weighted phase lag index (dwPLI) based connectivity of the prefrontal cortex (MNI: −20, 50, 40) in the theta band, which was selected in 15 % of SVMs. All other features were selected with a frequency of less than 10 %.

Thus, a multivariate machine learning approach could statistically significantly distinguish between chronic pain patients and healthy controls based on EEG measures of brain activity and connectivity. In particular, frontal phase-based connectivity at theta and gamma frequencies provided important information for the classification.

## Discussion

In the present study, we systematically exploited the potential of EEG to determine abnormalities of brain function in chronic pain. Defining such abnormalities promises to advance the understanding of the neural basis of chronic pain. Moreover, they might serve as a brain-based marker and novel treatment target of chronic pain. To this end, we analyzed resting-state EEG recordings of a large cohort of patients suffering from chronic pain and compared them to those of age- and gender-matched healthy controls. The analyses ranged from simple global measures of brain activity to sophisticated connectivity and network analyses in source space. All analyses were data-driven and rigorously corrected for multiple comparisons. To the very best of our knowledge, this approach represents the most comprehensive analysis of EEG data from one of the largest cohorts of chronic pain patients so far. The results show that global measures of brain activity and brain connectivity as measured by EEG did not differ between chronic pain patients and a healthy control group. However, our approach revealed a stronger phase-based connectivity at theta and gamma frequencies in the prefrontal cortex of chronic pain patients. Furthermore, we observed a global reorganization of brain networks in the gamma frequency band. Based on patterns of brain activity and connectivity, a multivariate machine learning approach could classify chronic pain patients and healthy controls with an accuracy of 57%.

Previous resting-state EEG studies investigating alterations in chronic pain patients mainly reported an increase in theta power together with a slowing of the global peak frequency compared to healthy controls [10; 65; 84; 88]. These findings have been related to the thalamocortical dysrhythmia model of chronic pain (TCD) [47; 48]. In this model, abnormal nociceptive input causes abnormal thalamic bursts at theta frequencies. These theta oscillations are transmitted to the cerebral cortex where they result in disinhibition of neighboring areas, abnormal oscillations at gamma frequencies and eventually in ongoing pain. This model is highly appealing, but evidence is still sparse. The present completely data-driven approach in a large cohort of chronic pain patients neither shows increased theta power nor a shift of global peak frequency and therefore does not directly support the TCD model of chronic pain.

The univariate comparisons of brain activity/connectivity between groups and the multivariate machine learning approach congruently indicated increased functional connectivity of the prefrontal cortex in chronic pain patients. These findings are in accordance with functional magnetic resonance imaging [5; 35; 80] and EEG [53] studies as well as with recent reviews and theories [4; 62; 69], which have shown that structural and functional alterations in the prefrontal cortex play an important role in chronic pain. A more precise localization of the connectivity increases in the prefrontal cortex is beyond the spatial resolution of EEG. Hence, it remains unclear how the present observations relate to the multitude of functions represented in the prefrontal cortex, which include motor, cognitive control, emotional, evaluative and modulatory functions [19; 42]. However, a role of the prefrontal cortex in chronic pain points to an important function of emotional-evaluative, motivational and decision-making circuits rather than sensory circuits in chronic pain [4; 62].

Our findings revealed that chronic pain is associated with focal increases of connectivity at gamma frequencies in the frontal cortex. These focal increases were associated with a global disturbance of brain network organization in the gamma band. The machine learning approach specifies that the focal frontal increase in gamma connectivity has the highest predictive value for distinguishing chronic pain patients from healthy controls. Mechanistically, gamma oscillations have been related to the activity of inhibitory parvalbumin-positive GABAergic interneurons [12]. In an animal model of chronic pain, these interneurons have been implicated in the modulation of pyramidal cell firing in the prefrontal cortex and pain behavior [95]. This link between GABAergic inhibition, gamma oscillations, prefrontal cortex activity and pain behavior is in accordance with the present observations. Functionally, gamma oscillations likely represent a basic feature of neuronal signaling and communication [22; 30; 90], which appears to be particularly related to the local processing and feedforward communication of currently important stimuli [22; 30; 59]. These concepts would be in line with an association of chronic pain with prefrontal gamma oscillations possibly signaling the emotional, motivational and evaluative aspects of pain.

Additionally, decreases of global efficiency and hub disruption index in the gamma band indicate that chronic pain is not only associated with focal changes of functional connectivity but also a global reorganization of brain networks. This is in accordance with recent fMRI studies which have shown global changes of functional connectivity at infra-slow frequencies below 0.1 Hz [50; 51]. The present findings complement these studies by showing that global changes of functional connectivity also occur in the gamma band at much higher frequencies between 60 and 100 Hz.

The present machine learning approach shows that applying an SVM classifier to resting-state EEG data allows to distinguish between chronic pain patients and healthy controls with 57% accuracy. A recent study which pursued a closely related EEG approach showed accuracies of > 90% for the classification of chronic pain patients vs. healthy controls [84], which were not achieved in the present study. The reasons for this disparity remain unclear as the available information on the previous approach does not allow for precise replication. Even though the accuracy of the present study is far from being sufficient for practical purposes, this result has important implications. First, it indicates that prefrontal synchrony at theta and gamma frequencies might play a role in the pathophysiology of chronic pain. Second, the present approach might represent a step further towards a much sought-after brain-based marker of pain [18; 79]. FMRI recordings have already shown that, in principle, it is possible to establish such a marker [50; 89]. The present approach complements these fMRI approaches by using EEG recordings. Third, abnormal patterns of EEG activity in chronic pain might represent potential targets for novel therapeutic strategies such as non-invasive brain stimulation techniques [60] and/or neurofeedback approaches [72]. In particular, the emerging transcranial alternating current stimulation technique [60] allows for the frequency-specific modulation of neuronal oscillations and synchrony and might, thus, represent a promising approach to modulate pain.

Several limitations of the present study need to be pointed out. First, abnormal oscillations and synchrony are observed in many neurological diseases [77; 78] and the specificity of the present results for chronic pain remains unclear. In particular, changes of prefrontal brain activity [61] and gamma synchrony [27] have also been observed in depression which frequently co-occurs with chronic pain [26]. However, changes of prefrontal theta and gamma synchrony were similarly found when patients with depression were excluded. Moreover, a potential lack of specificity does not necessarily limit the clinical usefulness and validity of a brain-based marker of chronic pain (see [18] for an in-depth discussion of criteria for brain-based markers of chronic pain). In fact, many well-established laboratory, electrophysiological and imaging tests yield results which are neither disease-specific nor symptom-specific, but are clinically highly useful. Second, the observed abnormalities of brain function were not significantly related to disease characteristics which suggests that they characterize the state of chronic pain *per se* rather than its quantitative characteristics. Third, field spread and/or muscle artifacts can cause spurious synchrony of EEG signals. A rigorous artifact correction procedure and analysis in source space are best practice to reduce these effects. Moreover, the present analyses are based on comparisons between groups and systematic differences of volume conduction between groups are unlikely. However, muscle artifacts and volume conduction remain an inherent and delicate confound of EEG signals. Fourth, drug effects cannot be ultimately ruled out. We excluded patients taking benzodiazepines, which have known effects on EEG signals [6]. However, in our representative cohort of chronic pain patients, most patients took non-opioid analgesics, opioids and/or antidepressants. To control for drug effects, we quantified medication and found no significant correlations between medication and the observed EEG effects. Therefore, it is unlikely but not impossible that our effects are solely driven by drug effects.

In conclusion, our data-driven, systematic and comprehensive analysis of EEG data from a large cohort of chronic pain patients shows that local and global measures of brain activity did not differ between chronic pain patients and a healthy control group. These negative findings might help to clarify inconsistencies in previous studies and guide future research. Moreover, our study reveals increased prefrontal synchrony together with global network reorganization at gamma frequencies in chronic pain which allows for differentiating chronic pain patients from healthy controls. These findings advance the understanding of the brain mechanisms of chronic pain. Beyond, the present observations might represent a step further towards a safe, low-cost, broadly available and potentially mobile brain-based marker of pain. However, substantial challenges concerning the accuracy, specificity and validity of such a marker remain to be overcome. Finally, the findings might open new therapeutic perspectives by revealing a potential target for novel non-invasive brain stimulation and neurofeedback strategies.

## Acknowledgments

The authors kindly thank all patients and healthy participants for their participation in the study. Special thanks to Christine Berger and Vesna Dolanjski for support with patients and Marlene Försterling for help with data acquisition. We thank Ulrike Bingel and Valentin Riedl for helpful comments on the manuscript. The study was supported by the Deutsche Forschungsgemeinschaft (PL 321/10-1, PL 321/11-1, PL 321/10-2, PL 321/11-2), the Bavarian State Ministry of Education, Science and the Arts, and the Wellcome Trust (098433). The authors declare no competing financial interests.

